# Characterization of Arabian Peninsula whole exomes: exploring high inbreeding features

**DOI:** 10.1101/2022.02.22.481461

**Authors:** Joana C. Ferreira, Farida Alshamali, Luisa Pereira, Veronica Fernandes

## Abstract

The exome (WES) capture enriched for UTRs on 90 Arabian Peninsula (AP) populations contributed nearly 20,000 new variants from a total over 145,000 total variants. Almost half of these variants were in UTR3, reflecting the low effort we have dedicated to cataloguing these regions, which can bear an important proportion of functional variants, as being discovered in genome-wide association studies. By applying several pathogenic predicting tools, we have demonstrated the high burden in potentially deleterious variants (especially in nonsynonymous and UTR variants located in genes that have been associated mainly with neurologic disease and congenital malformations) contained in AP WES, and the burden was as high as the consanguinity level (inferred as sum of runs of homozygosity, SROH) increased. Arabians had twice SROH values in relation to Europeans and East Asians, and within AP, Saudi Arabia had the highest values and Oman the lowest. We must pursuit cataloguing diversity in populations with high consanguinity, as the potentially pathogenic variants are not eliminated by genetic drift as much as in less consanguineous populations.

## Introduction

The technological developments introduced with the next-generation sequencing methodology are allowing the high throughput characterization of millions of polymorphisms, enhancing the knowledge on genetic diversity between and within populations, and enabling new lens to explore the genetic basis of diseases [1!,2]. Directed sequencing of all exons (known as whole exome sequencing, WES) is being increasingly used in clinical genetics [3], as these regions are probable locations of candidate alleles conferring susceptibility to monogenic and complex diseases. International consortia have been collecting big exome data and making them publicly available for mining of variants [4,5], such as the Exome Sequencing Project (ESP) Exome Variant Server (EVS) [6] representing more than 200,000 individuals from multiple ESP cohorts, and the Exome Aggregation Consortium (ExAC) database [7] spanning 60,706 unrelated individuals sequenced as part of various disease-specific and population genetic studies. The ExAC creators launched afterwards the Genome Aggregation Database (gnomAD), which englobes currently 125,748 WES and 15,708 WGS [8]. This big data is allowing to predict the functional effects of variants on diverse traits in different populations, and guiding inference of disease risk at an individual basis. However, WES catalogues are still extremely biased in terms of ancestry, being largely based on data from European and East Asian populations [4,9], which limits the transferability of findings to other population groups [10].

A region particularly understudied in large-scale sequencing projects is the Middle East and more specifically the Arabian Peninsula (AP). AP populations are amongst the most non-African ancestral populations in the globe, as this region was the first outpost of the successful out-of-Africa (OOA) migration, at around 60 thousand years ago (ka) [11,12]. Concordantly, signatures of ancient ancestry were detected in extant AP populations, especially in the Arabo-Persian Gulf region: (1) as relic maternal lineages from the west-Eurasian haplogroups N1, N2 and X, branching directly from the root of macro-haplogroup N (Fernandes et al. 2012); (2) in the autosomal genome, the component known as basal Eurasian, identifiable when ancient DNA information is integrated in the analysis (Ferreira et al. 2021). There were clear continuums throughout time linking western Arabia with the Levant and Africa, and eastern Arabia with Iran and the Caucasus, making AP an important migration nexus between the main human population groups [13,14]. This high admixture enriched the Arabian genomes with non-autochthonous selected variants, mainly for malaria protection (higher in the west), lactose tolerance (European/South Asian-derived allele in the eastern AP) and other immune system defences (throughout the Peninsula) [13,15].

Despite the high ancestry admixture in AP, Arabs have traditionally a high level of consanguinity, reaching an overall prevalence of 56% in Saudi Arabia, from which 33.6% are first-cousin consanguineous marriages [16]. In addition to a high consanguinity rate, Arabs are also characterized by large family sizes and advanced paternal and/or maternal age [17], increasing the risk of congenital anomalies in their offspring [16]. These factors are associated with a high number of congenital and genetic disorders [17], such as impaired hearing (3.5 times higher in consanguineous than in non-consanguineous mating) [16] and Down’s syndrome (Arab countries exceeds the 1.2-1.7 per 1000 typical for industrialised countries; [17,18]). Other complex disorders with a genetic component are also common throughout the Arab world, including haemoglobinopathies, glucose-6-phosphate dehydrogenase deficiency and metabolic diseases (obesity, type 2 diabetes and dyslipidemia), and all have been associated with the high level of consanguinity in this region [16]. The reason for this is that high level of consanguinity increases frequencies of rare variants and extends stretches of homozygous chromosomal fragments (long runs of homozygosity across all sites, ROH) [19]. Thus, the study of ROH length and burden of deleterious variation can provide important insights into the human demographic history and clinical applications [20,21]. Most of the WES available for Arabians were obtained in cohorts of patients [22-25]. Fewer WES are available for the general population: one obtained in Qatari populations identified eight hematologic variants, five metabolic, four eye-related, three inflammatory, three cardiovascular and three neurologic as the most common disease-related variants in those consanguineous cohorts [26]; an important cohort, the Greater Middle East (GME) Variome Project containing 2,497 individuals from 19 Arab and non-Arab Muslim countries [27] detected large and rare homozygous blocks, compatible with recent consanguineous matings, rendering easy to identify genes harbouring putatively high-impact homozygous variants.

As the effective size of the WES catalogue for AP general population remains low, in this work we performed a characterization of Arabian WES by randomly selecting 25 individuals from each AP country (Saudi Arabia, Yemen, United Arab Emirates and Oman). We enriched our WES with the sequencing of the untranslated regions 5 and 3 (UTR5 and UTR3), usually not included in these panels, as these regions play a role in the modulation of mRNA transcription, secondary structure, stability, localization, translation, and access to regulators like microRNAs and RNA-binding proteins [28]. We aimed to identify new and rare variants (especially in UTRs), to infer their possible functional impact, and to relate them with the consanguinity levels.

## Materials and methods

### Sample collection and WES capture

We conducted WES of 94 individuals from AP (23 from Saudi Arabia, 24 from Yemen, 25 from Oman and 22 from UAE). These samples were part of a larger cohort of 420 Dubai residents who were born across the AP. As this cohort was previously analysed with the Illumina Human Omni Express Bead Chip containing 741,000 SNPs [13,29], it allowed us to selected the 94 individuals for WES based on the genomic information that they were non-related and non-recent migrants from sub-Saharan Africa. This study obtained the ethical approval from the Ethics Committee of the University of Porto, Portugal (17/CEUP/2012).

WES was performed at two companies: Macrogen Inc. (Seoul, South Korea) for 50 samples with a 100x average depth coverage; and STAB VIDA (Caparica, Portugal) for 44 samples with a 30x average depth coverage. The WES were captured using the SureSelect Human All Exon V5 + UTRs (50 Mb) target enrichment kit. Sequencing was performed with 2 × 100 bp paired end reads on Illumina HiSeq platform (Illumina, San Diego, CA, USA) according to manufacturer’s protocol.

In order to have comparable values for other worldwide populations, the 1000 Genomes [30] WGS from African, European and East Asian populations (90 individuals randomly selected from each of these regions; Supplementary Table 1) were extracted, and variants located in the genomic regions covered by our WES (including regulatory untranslated regions UTRs) were considered for further analyses in this manuscript.

### Variant calling, filtering and annotation

Paired-end reads were trimmed using the trimmomatic tool and aligned to human reference genome NCBI Build 37 using the Burrows-Wheeler Aligner (BWA v0.7.16a) algorithm. The paired-read alignments were sorted and stored in BAM format using samtools (v1.5). All the samples passed the quality control performed with FastQC and Alfred tools. Duplicates were marked and eliminated with Picard (v1.139), and local re-alignment and Base Quality Score Recalibration were carried out using the Genome Analysis Toolkit (GATK, v4.2.0.0, https://software.broadinstitute.org/gatk/).

Variants were called using the Genome Analysis Toolkit (GATK v4.2.0.0) Haplotypecaller. A minimum value of 10% was set for missing genotype, leading to exclusion of three samples from further analyses. Variants were initially filtered to have a minimum depth of 7, Phred quality score >30, and genotype quality score >20, using the bcftools (v2.26.0). In the case of heterozygous callings for which the ratio of the less covered allele (reference or derived) over the total calls was <25%, the genotyping was corrected to homozygous of the most frequent allele, by using an in-house script. Finally, we excluded from the analysis multi-allelic variants, indels and positions for which more than 5% of the genotypes were missing. A principal component analysis (PCA) was performed using the SmartPCA tool from the EIGENSTRAT software package (v6.1.3), to identify potential batch effects between laboratories, or outliers. After the PCA, one sample was eliminated as it was a clear outlier. The final dataset contained 90 samples (23 from Saudi Arabia, 24 from Yemen, 24 from Oman and 19 from UAE). To further assess the quality of the calling, we evaluated in the bcftools stats the aggregate transition-to-transversion (Ti/Tv) ratio for all variants and exonic variants.

The ANNOVAR tool (version available in 2019-10-24; [31]) was used for the functional annotation of the called variants, and to verify if they were previously described in the following publicly available databases (downloaded on 10^th^ September 2021): WGS group, which included the GnomAD_WGS V2.0.1 (11st March 2017), dbSNP_142 (28^th^ December 2014), Haplotype Reference Consortium (HRC; 3^rd^ December 2015), and 1000 Genomes (24^th^ August 2015); WES group, consisting in GnomAD_exome V2.0.1 (11st March 2017), NHLBI-ESP (esp6500siv2_all; 22^nd^ December 2014) and ExAC (29^th^ November 2015); ClinVar (clinvar_20160302; 3^rd^ October 2017) and dbNSFP3.0a group; and finally, Greater Middle East group (GME; 24^th^ October 2016).

### Functional constraint analysis

The online CADD tool (v1.6; https://cadd.gs.washington.edu/snv) was used to evaluate the evolutionary conservation of AP (and comparable 1000 Genomes datasets) variants, through several metrics. The first type of metrics considered only alignment-based conservation values: (1) GERP or Genomic Evolutionary Rate Profiling [32], which evaluates non-neutral rates of substitution from multiple mammalian species alignments, and categorizes mutations by their predicted deleterious effect in neutral (−2< GERP < 2), slightly deleterious (2 <GERP< 4), moderate (4 < GERP< 6), and extremely deleterious (GERP> 6) groups (categories according to [33,34]); (2) Phylop [35], also based on the alignment of mammalian species, for which a negative value indicates faster-than expected evolution, while positive values imply conservation.

Secondly, the metrics SIFT (Sorting Intolerant From Tolerant; [36]) and PolyPhen [37] that predict whether nonsynonymous substitutions are likely to have a deleterious effect on the protein function were investigated. A nonsynonymous variant with a SIFT score < 0.05 will be classified as ‘deleterious’ while others are called ‘tolerated’ (benign) [38]. In contrast, PolyPhen2 calculates the probability that a given variant will be ‘benign’ for scores less than or equal to 0.446, ‘possibly damaging’ for scores greater than 0.446 and less than or equal to 0.908, and ‘probably damaging’ for scores greater than 0.908.

Finally, the integrative “scaled C-score” was considered. This score provides a ranking of variants more likely to be deleterious by integrating multiple annotations including conservation and functional information into one metric [39]. We applied a cutoff of 20, below which the variants were classified as benign and otherwise harmful, as suggested by the authors [39].

### Inbreeding features

To infer the degree of relatedness between the 90 AP WES we used the KING tool [40] by estimating kinship coefficients and inferring IBD segments for all pairwise relationships. Unrelated pairs can be precisely separated from close relatives with accuracy up 4th-degree.

To assess the individual runs of homozygosity (ROHs), we estimated ROHs using the PLINK tool (version 1.9) following these published [41] parameters: a size threshold (kb) to call an ROH (homozyg-kb) of 1000 kb; a SNP number threshold to call an ROH (Homozyg-snp) of 10 SNPs; a sliding window size in SNPs (Homozyg-window-snp) of 20 SNPs; allowing 5 missing SNPs (Homozyg-window-missing); with a proportion of homozygous windows threshold (Homozyg-window-threshold) of 0.05; a minimum SNP density of 200 kb to call an ROH (Homozyg-density); allowing a maximum gap (Homozyg-gap) of 4000 kb; and allowing only 1 heterozygous SNP (Homozyg-window-het). For each AP individual included in the analysis, the sum of the length of ROHs (SROH) was calculated. These AP SROHs were used for testing the linear regression with the burden of predicted pathogenic non-synonymous variants (inferred for both SIFT and PolyPhen algorithms), and a f-statistic test was applied.

### Mining of diseases associated with highlighted genes

Information for disease associated with genes for which AP individuals presented predicted pathogenic non-synonymous and conserved UTR variants was collected from the OMIM (https://www.omim.org/) database. Then, these diseases were mined in the MalaCards: The human disease database (https://www.malacards.org/) for classification in broad categories. In cases of disease affecting several organs, decisions were made for inclusion in one category instead of another (Supplementary Table 2).

### Graphs and statistical tests

The plots were built with the venn [42,43] and ggplot2 package [44] in R [45]. The calculations for those plots were performed through in-house R-scripts.

## Results

### Diversity of the AP WES (enriched for UTRs)

The 90 AP WES presented 145,630 variants, around half the level of diversity displayed by the European (270,913 variants) and East Asian (254,527 variants) regions, and one third by sub-Saharan Africans (443,767 variants) (1000 Genomes populations, for an equal number of individuals and matching genomic segments). The transition-to-transversion ratio (Ti/Tv) was 2.59 for the entire dataset, and 3.23 when considering only the exonic variants as this latter value is more comparable with other high quality WES datasets not enriched for UTRs [1,2].

Of the total 145,630 variants, 19,701 (13.5%) were new when compared against several public databases (Figure 1A). Among the known variants (Figure 1B; Supplementary Table 3), identical proportions (around a quarter) were shared between nonsynonymous, intronic and UTR3 classes, followed by synonymous (∼16%) class, and rare proportions of all remaining classes (nonsense, UTR5, ncRNA, upstream/downstream, intergenic and splicing). Interestingly, for the new variants (Figure 1C), almost half of these were located in UTR3, followed still by 29% of intronic, 14% of synonymous and rare instances of the other classes of new variants. It is not surprising the high proportion of new variants in UTR3, and that the value of new variants in UTR5 is also twice the value for this class in known variants, as UTRs are rarely covered in WES screenings. We detected very few new nonsynonymous variants (57 - <1% of total new variants), showing that saturation for this class of variants is almost reached for populations of mainly Eurasian ancestry.

**Figure 1:**
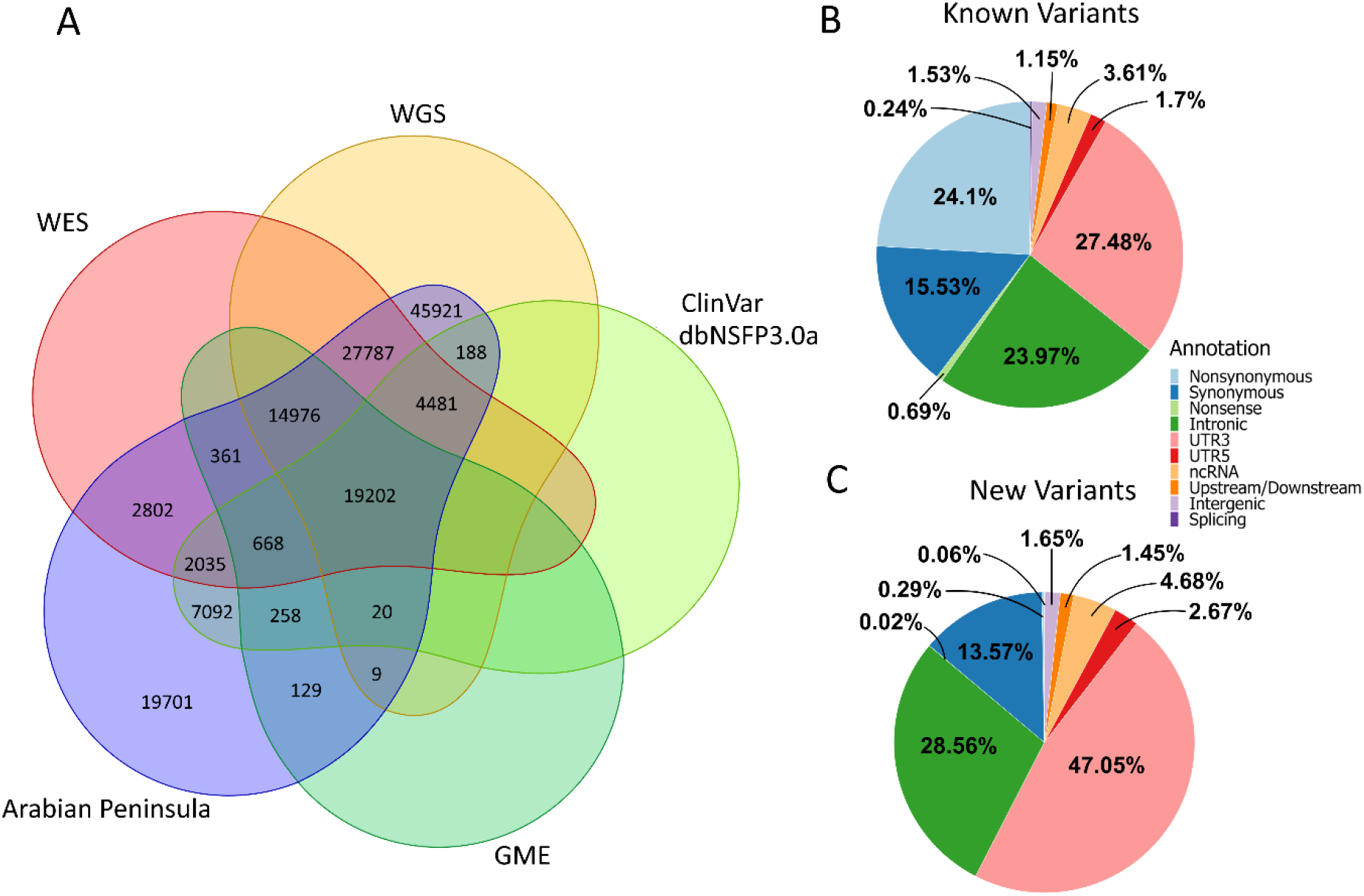
The global diversity observed in the 90 AP WES. **A**. Venn diagram illustrating new and shared variants between AP WES and publicly available databases (as referred in the material and methods section). **B**. Classes of known variants observed in the AP WES. **C**. Classes of new variants observed in the AP WES.

Interestingly, a substantial amount of AP variants (7092) was only shared with clinically/functionally focused databases, namely ClinVar [46] and dbNSFP3.0a [47]. A careful inspection of these variants revealed they were mostly included in the dbNSFP3.0a dataset (88.8% non-synonymous, 6.78% non-sense and 2.19% splicing variants), which prioritizes these type of variants from UK10K and ExAC datasets. These variants are of possible functional relevance, as we will see in detail in the next section.

### Functional constraint features of AP WES

The conservation score GERP (Figure 2A and B) allowed to confirm that AP had 83.9% of variants classified either as neutral (66.2% for GERP<2) or as slightly deleterious (17.7% for 2 >GERP> 4). These variants were broadly distributed by class of variants, testifying that most of the genome of AP is functionally neutral as expected. Comparatively, in the two most deleterious categories (4 > GERP> 6 and 4 > GERP> 6) there was an increase of nonsense, nonsynonymous and splicing variants, and a decrease of UTR5 and UTR3. The results from the Phylop algorithm, another conservation score, were consistent with GERP (Supplementary Figure 1). When comparing with the other worldwide regions (Figure 2C), AP displayed the higher amount of moderate and extremely deleterious variants (15.31% and 0.81% respectively; compensated by the lower amount of neutral variants), followed *ex aequo* by Europe (11.22% and 0.37% respectively) and East Asia (11.49% and 0.39% respectively), and then Africa (10.18% and 0.30% respectively) with the lower values.

**Figure 2:**
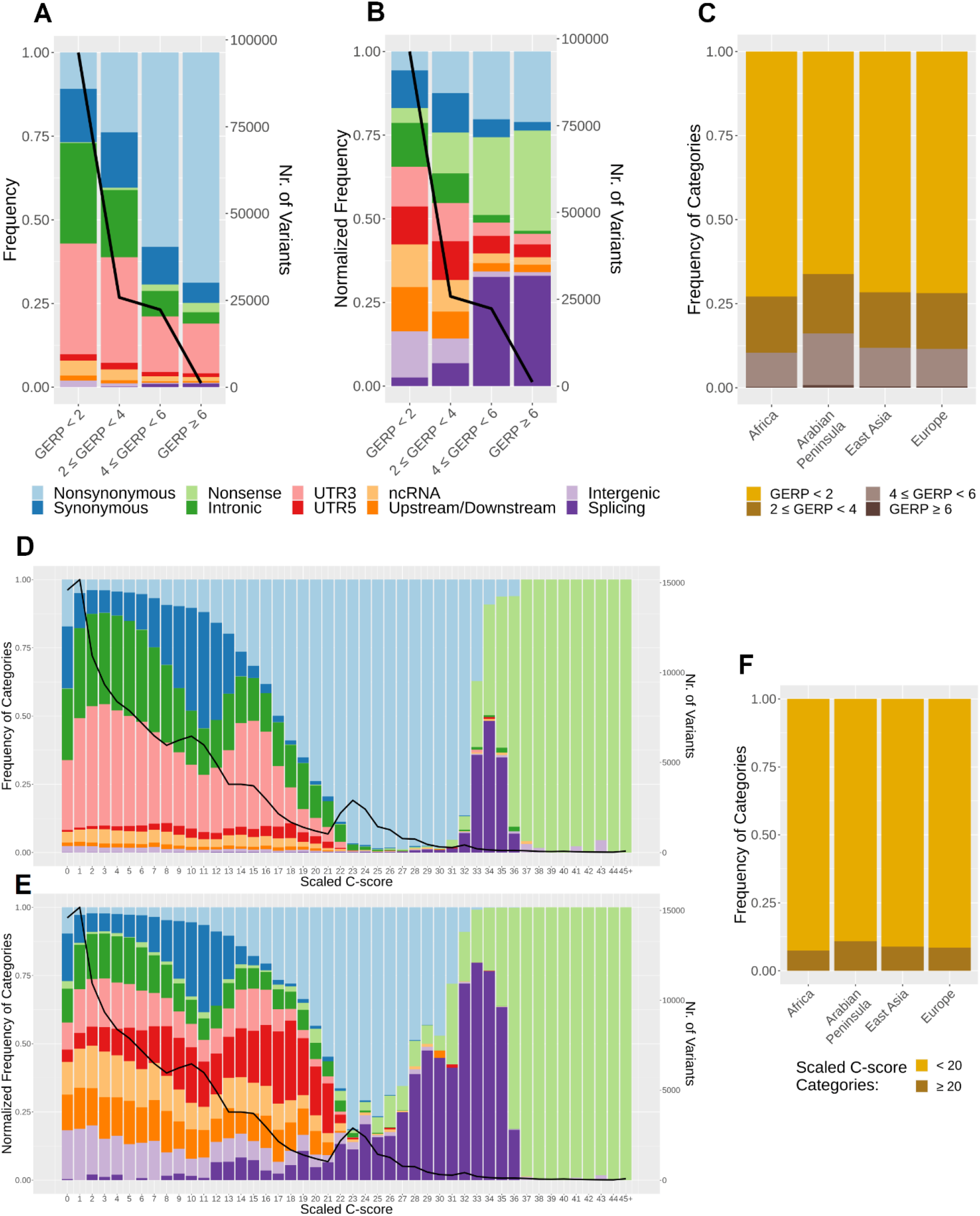
Results for the conservation GERP and the integrative scaled C scores. **A**. Proportion of AP variants from each class of variants in each GERP category. The line represents the total number of variants in each score category (same meaning for B, D and E). **B**. Proportion of AP variants after normalizing by the total number of variants in each class, observed in that GERP category. **C**. Proportion of variants from each GERP category in AP and other worldwide regions. GERP values were classified into four groups: neutral (2< GERP), slightly deleterious (2 <=GERP< 4), moderate (4 <= GERP< 6), and extremely deleterious (GERP> 6). **D**. Proportion of AP variants from each class of variants along the scaled C-scores. **E**. Proportion of AP variants after normalizing by the total number of variants in each class, along the scaled C-scores. **F**. Proportion of variants with scaled C-scores below and above 20 (benign and harmful variants, respectively) in AP and other worldwide regions.

A similar pattern was observed for the integrative scaled C-scores, but now with better resolution (Figure 2D and E). Nonsense variants are the most extreme in the harmful scale, preceded by the splicing variants. For non-synonymous variants, a high percentage of them (45,89%) displayed values equal or higher to 20, so more prone to have functional impact. In the other extreme, variants that are of lower functional impact, were the synonymous, intronic, UTRs, and the rarer ncRNA, upstream/downstream and intergenic variants. Comparing with the other geographical regions (Figure 2F), again AP had a higher proportion of potentially harmful variants when inferred through the scaled C-scores (10.86%) than the rest of the globe (East Asia – 8.89%; Europe – 8.52%; Africa – 7.38%).

Focusing on the predicted pathogenicity for AP nonsynonymous variants, provided by SIFT and PolyPhen metrics (Figure 3A), a proportion of 15.56% (4,673 out of 30,033 nonsynonymous variants for which values for both metrics were available) were inferred as pathogenic by SIFT and PolyPhen (“deleterious” and “probably damaging”, respectively). These variants were distributed in 3,285 genes, and 756 of these genes have been associated with diseases of various types (Figure 3B), namely (in decreasing importance) neurologic, congenital malformation, metabolic, ear and eye, immune and infection-related, oncologic, cardiovascular and blood. Only six of these inferred as pathogenic nonsynonymous variants were new, and they were located in the genes: *CYFIP1*, associated with congenital malformation and neurologic diseases; *ITGA10* and *PIAS3*, associated with oncologic, and ear and eye diseases; *GNG14* involved in signalling pathways; *C1GALT1C1L* that enables the activity of a galactosyltransferase whose deficit is associated with congenital malformation; and *FAM240A* that is mainly expressed in brain.

**Figure 3:**
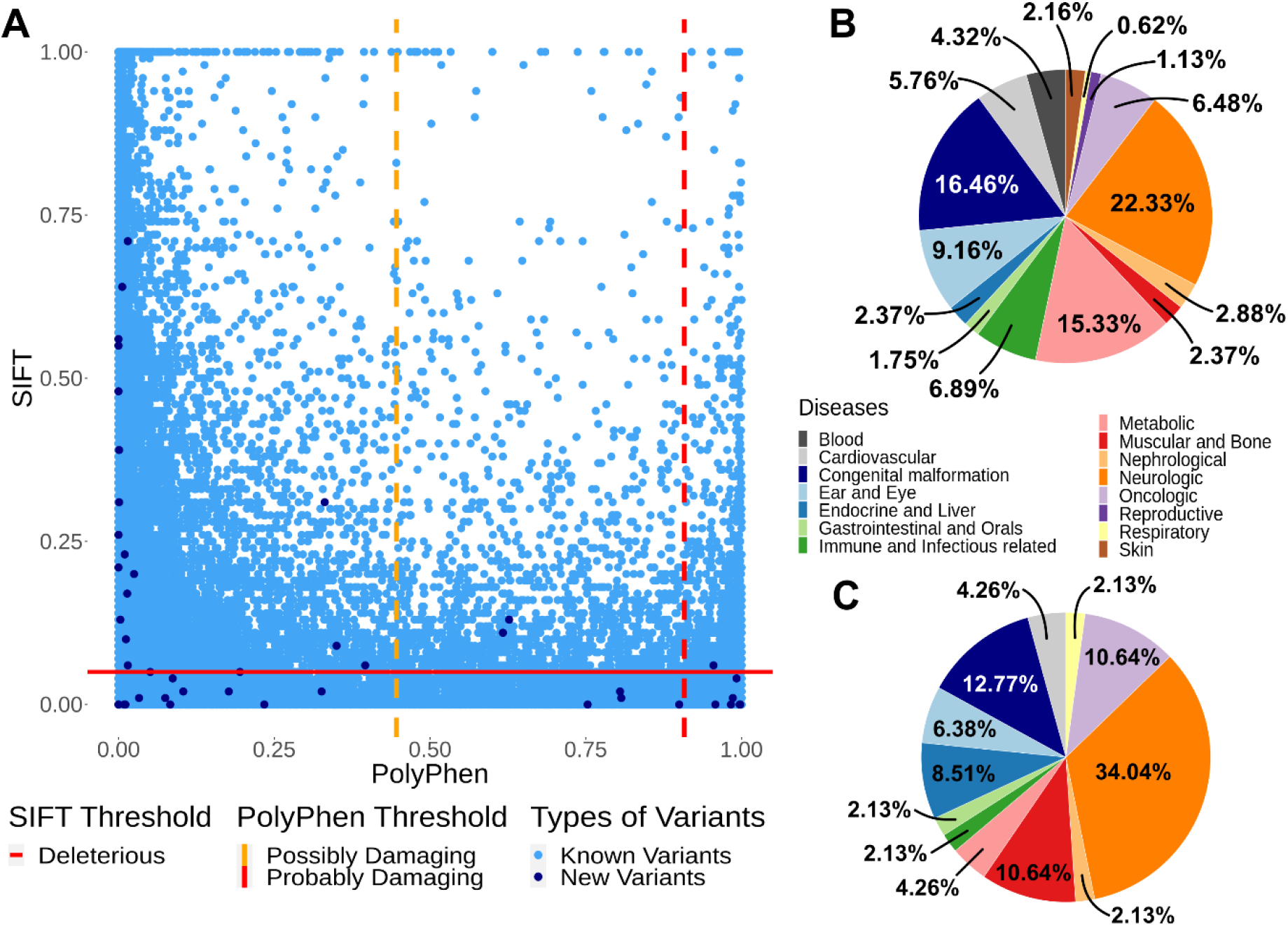
Pathogenic predictions. **A**. SIFT scores versus PolyPhen2 scores. The known variants are represented in light blue dots while new variants are represented as dark blue dots. The yellow dashed line correspond to the thresholds for predicting 0.446<Possibly Damaging<0.908 for PolyPhen2. The red dashed lines correspond to the thresholds for predicting deleterious variants: <0.05 for SIFT and Probably Damaging >0.908 for PolyPhen2. **B**. Diseases associated with 756 of the genes bearing predicted pathogenic (by both SIFT and PolyPhen) non-synonymous variants in AP, according to OMIM (classification of disease based on MalaCards database; Supplementary Table 4). **C**. Diseases associated with 31 of the genes bearing non-conserved (C-score above 20) UTR3 and UTR5 variants in AP, according to OMIM (classification of disease based on MalaCards database; Supplementary Table 4).

Our AP WES had the special feature of being enriched for the UTR regions, providing additional information for these parts of the genome that, although untranslated, can be pivotal in the regulation of gene expression. As we have referred before, most of UTR variants are non-conserved, but focusing on the proportion of UTRs with C-scores above 20, 1.76% of UTR5 and 0.31% of UTR3 variants (47 out of 2671 UTR5 and 138 out of 43,868 UTR3) can be potentially harmful variants. These variants were distributed in 116 genes, and 31 of these genes have been associated with various types of diseases (Figure 3C), with the high percentage of them classified as neurologic diseases (34.04%) and congenital malformation (12.77%), compatible with the results observed for the inferred pathogenic nonsynonymous variants.

### Inbreeding features inferred from the AP WES

The inference of relatedness between the 90 AP WES, based on the genomic kinship coefficient, revealed that all the individuals were classified as unrelated (up 4th-degree), as expected for a cohort representative of the general population. We then estimated ROHs longer than 1Mb in AP and other worldwide populations, in order to compare inbreeding features. In concordance with the known higher consanguinity of AP populations, there were more ROHs of all sizes in Arabia than in the other regions of the globe (Figure 4A), mainly for the ROH size category 2-4Mb. The largest ROHs detected in each AP population were: Oman-70.4Mb; Yemen-59.8Mb; Saudi Arabia-41.2Mb; and UAE-26Mb.

**Figure 4:**
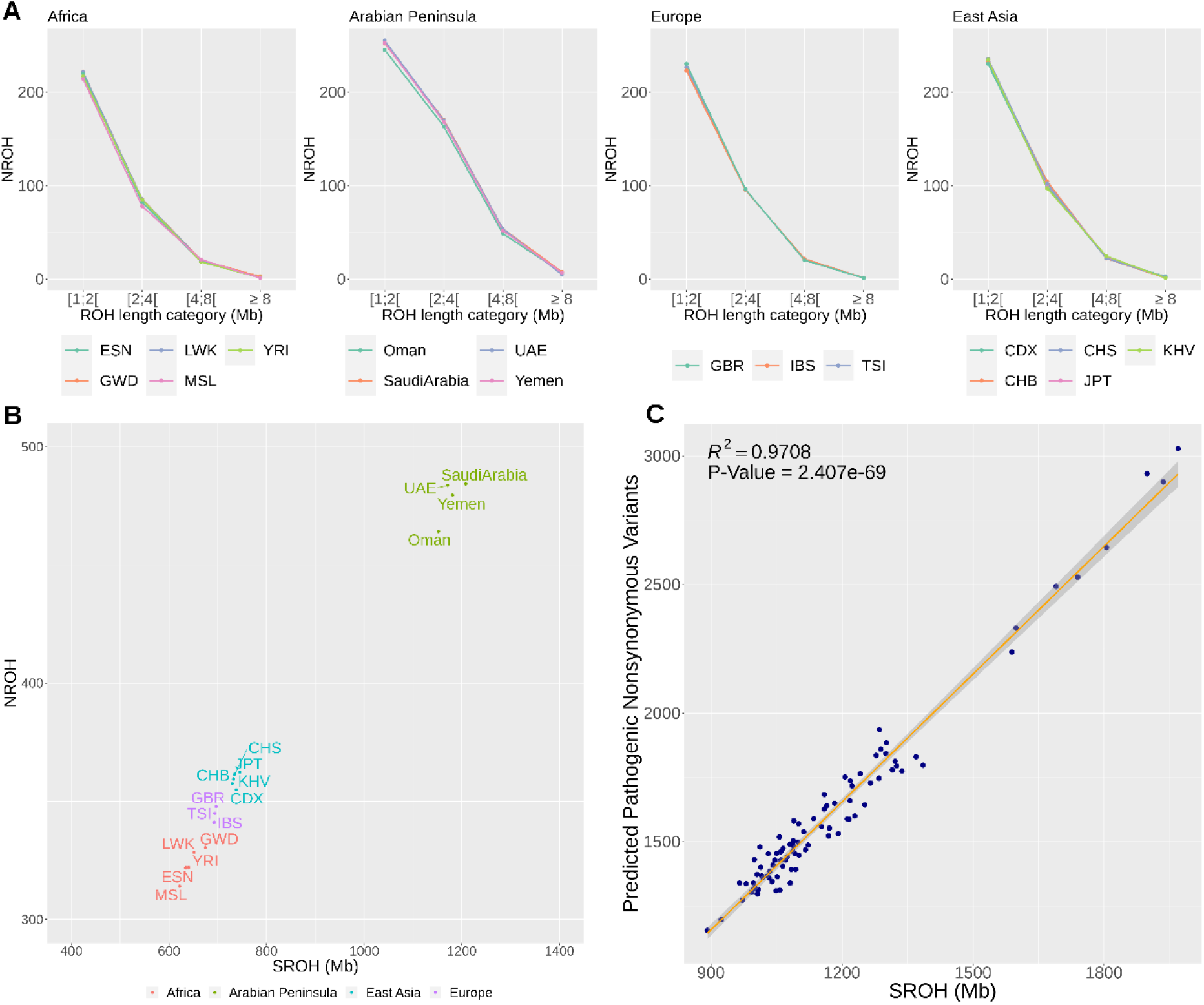
Inbreeding features. **A**. Mean number of ROHs, NROHs≥1 Mb per population in different length categories, for AP and other globe regions. **B**. NROHs versus the mean sum ROH length (SROH) in each region: Africa in red (ESN-Esan in Nigeria, GWD-Mandinka in Gambia, LWK-Luhya in Kenya, MSL-Mende in Sierra Leone, and YRI-Yoruba in Nigeria); AP in green; East Asia in blue (CDX-Dai in China, CHB-Han in China, CHS-Han in South China, JPT-Japanese in Japan, and KHV-Kinh in Vietnam), and Europe in purple (GBR-British in UK, IBS-Iberians in Spain, and TSI-Toscani in Italia). **C**. Linear regression between SROH and predicted pathogenic non-synonymous variants (for both SIFT and PolyPhen algorithms) in individuals from AP.

Not only AP individuals had larger ROHs, but also the total sum of ROHs (SROHs) per AP genome was in mean twice the values observed in the individuals from other worldwide populations (Figure 4B): 1200Mb in AP; 720Mb in East Asians, 700Mb in Europeans; and 650Mb in Africans. Within AP, the highest values were observed in Saudi Arabia and the lowest in Oman. In terms of SROH distribution within each population group, there was a high heterogeneity between values from AP individuals (Supplementary Figure 2), while individuals from other regions were more homogeneous in this metric. This highest SRHOs values in each AP country were: Saudi Arabia-1969Mb; Yemen-1781Mb; UAE-1171Mb; and Oman-1152MB.

Interestingly, within AP there is a very significant positive linear regression (r^2^=0.9712; p-value=1.315e^-69^) between SROHs and the burden of predicted pathogenic non-synonymous variants (inferred for both SIFT and PolyPhen algorithms; Figure 4C).

## Discussion

In the present study, we have performed a WES exome capture enriched for UTRs on AP populations, still poorly represented on international WES catalogues. We were able to identify 19,701 new variants (13.5% of the total 145,630), that were not catalogued in other public databases, and almost half of these were located in UTR3. We further confirmed that 185 of the UTR variants could be deleterious (inference based on the scaled C-score), being located in genes associated with various types of diseases, especially neurologic and congenital malformation, testifying the importance of screening UTR variants [28,48]. Our results match a previous report [28] focused on the UTR variants from the NHGRI GWAS Catalog [49], which were shown to be mostly associated with immunological, neoplastic and neurological pathologies. As the fine-mapping of UTR regions, especially in terms of their functional impact, is lacking behind the coding regions, there are currently no better pathogenic inferences for these variants than metrics based on conservation. As we improve our knowledge on the functional impact of UTRs, we will refine inferences about their role in diseases. These UTR variants add up to the substantial proportion in the AP WES of 4,673 nonsynonymous variants (15.56% out of 30,033 total) inferred as pathogenic by SIFT and PolyPhen metrics, and that were distributed in 3,285 genes, 23% of which have been associated also with neurologic and congenital malformation diseases, as well as other complex disorders.

We have demonstrated that this high burden on pathogenic variants in a relatively low diverse WES cohort (half the level of European and East Asian, and one third of sub-Saharan cohorts) can be explained by the demonstrated considerably high proportion of ROHs due to consanguinity practices. It is known that the strong bottleneck in the OOA dispersal led to a higher burden of pathogenic variants in Europeans and East Asians relative to sub-Saharan Africans [34], but the values for AP are impressive. The low diversity in the AP WES seems contra-intuitive to the identified high admixture in AP populations of sub-Saharan African and South Asian ancestries [13,14,29], but the high consanguinity is strong enough to opose the enrichment with diversity from other population groups. We must reinforce that we were careful in pre-selecting individuals with a main Arabian/NearEast background for WES screening, avoiding Arabian individuals with recent events of admixture. So, values of SROHs would still be more heterogeneous for AP populations if we had included these Arabians with more recent events of admixture. As it is, the main Arabian/NearEast background WES studied here contained several large and homozygous blocks, especially so in Saudi Arabia, where first-cousin marriages are common [16]. Two Saudi Arabian genomes had the highest sum of ROHs, as high as 2,000Mb. For contextualisation, the diploid human genome has 6,200Mb, so those Saudi individuals had ROHs across 32% of their genomes. In mean, the Arabian individuals had ROHs across 19% of their genomes, against 12% in East Asians, 11% in Europeans and 10% in sub-Saharan Africans. And as we are sequencing coding and regulatory regions of the genome, it is not surprising the high amount of predicted pathogenic variants we found.

Our findings highlight the importance of continuing to catalogue WES in general population cohorts, and in regions of the globe poorly represented in international consortia. The gains definitely pay off the efforts. A contribution of near 20,000 new variants from sequencing around 2% of the genome in 90 AP individuals is substantial. The pursuit of this cataloguing in populations with high consanguinity is advisable. As we have seen here, the higher the consanguinity the higher the burden of potentially pathogenic variants. These variants are harder to detect in less consanguineous populations due to the effect of genetic drift, which is opposed by consanguinity. In conclusion, we must enrich WES catalogues in ethnic groups and in populations with diverse breeding features, in order to increase the power to robustly identify disease-associated variants in the human species.

## Supporting information

https://cloud.i3s.up.pt/index.php/s/D6ApW4jjYEm7YtJ

## Acknowledgments

This work was financed by FEDER-Fundo Europeu de Desenvolvimento Regional funds through COMPETE 2020-Operacional Programme for Competitiveness and Internationalisation (POCI), Portugal 2020, by Portuguese funds through FCT-Fundação para a Ciência e a Tecnologia, Ministério da Ciência, Tecnologia e Inovação in the framework of the project “Biomedical anthropological study in Arabian Peninsula based on high throughput genomics” (POCI-01-0145-FEDER-016609). VF has a postdoc grant through FCT (SFRH/BPD/114927/2016). i3S is financed by FEDER-COMPETE 2020, Portugal 2020 and by Portuguese funds through FCT in the framework of the project ‘Institute for Research and Innovation in Health Sciences’ (POCI-01-0145-FEDER-007274).

## Data Availability

The WES from the individuals analyzed in this study can be accessed from the EGA repository (European Genome-Phenome Archive; accession number: EGA00000000000).

