## Supplementary material for "Characterization of Arabian Peninsula whole exomes: exploring high inbreeding features": https://cloud.i3s.up.pt/index.php/s/D6ApW4jjYEm7YtJ

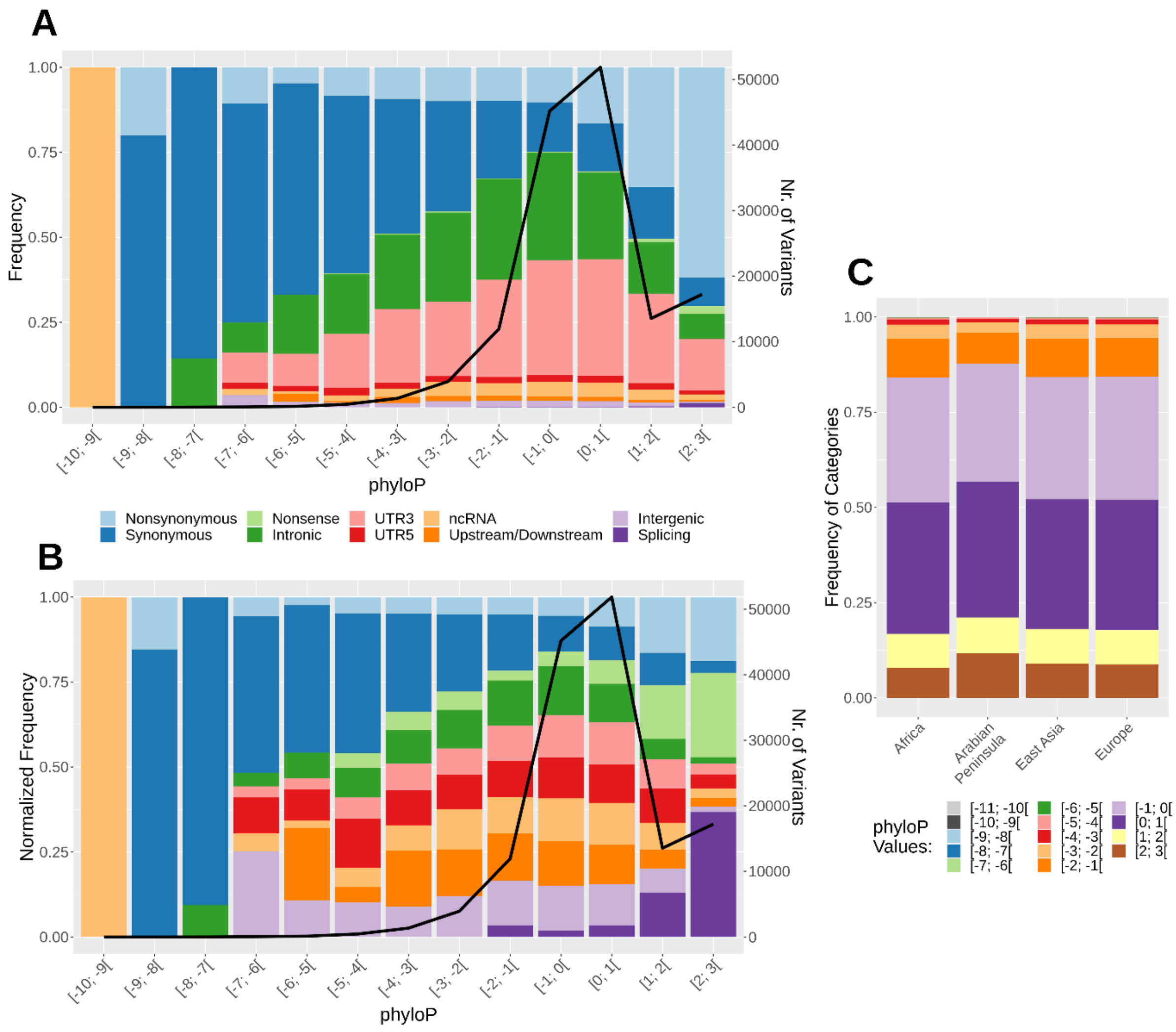


**Supplementary Figure 1.** Results for the conservation Phylop based on the alignment of mammalian species. A. Proportion of AP variants from each class of variants in each Phylop category. The line represents the total number of variants in each score category (same meaning for B). B. Proportion of AP variants after normalizing by the total number of variants in each class, observed in that Phylop category. C. Proportion of variants from each Phylop category in AP and other worldwide regions. Phylop values were classified into intervals of one and negative values indicates faster-than expected evolution, while positive values imply conservation.

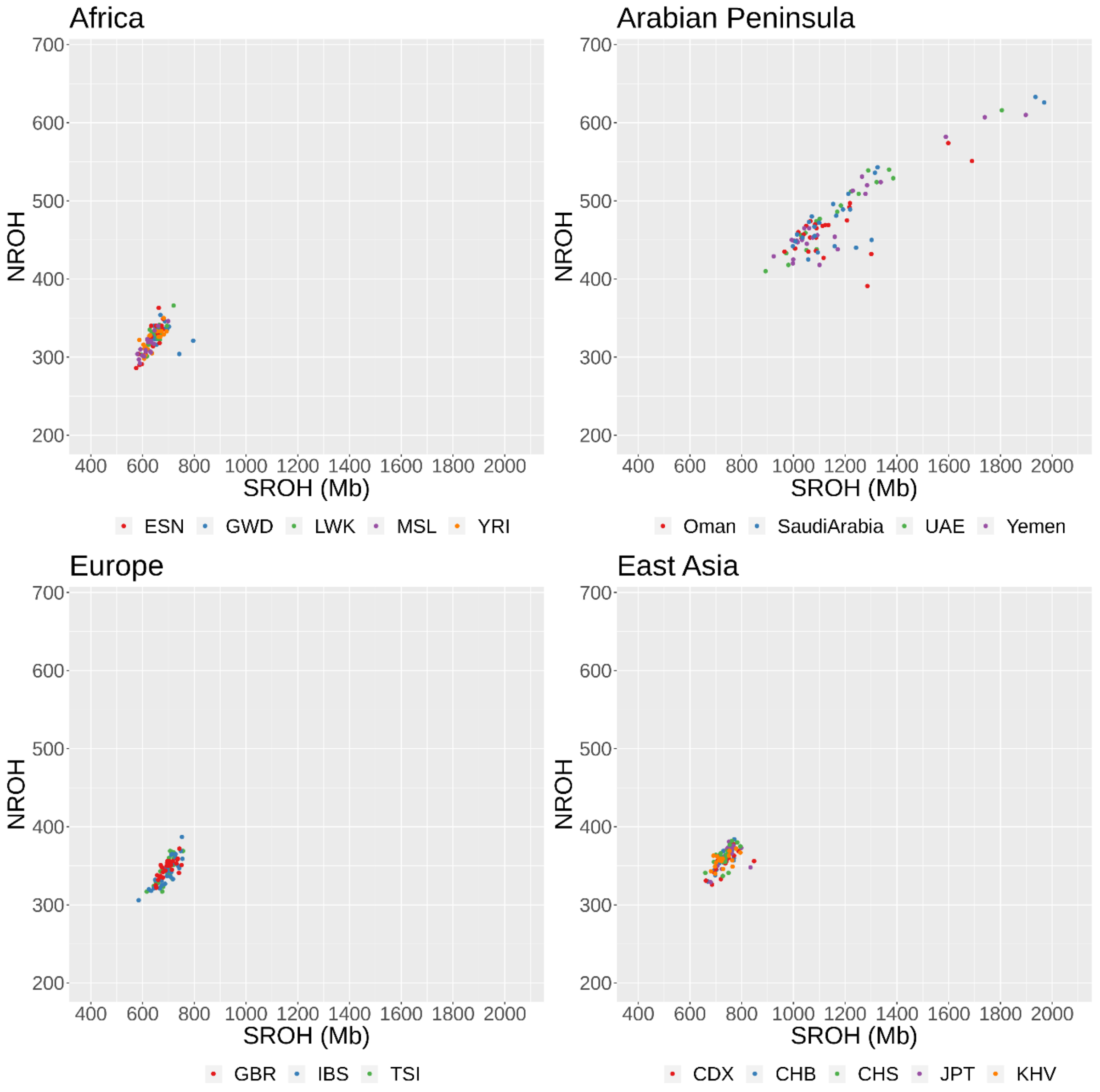


**Supplementary Figure 2.** Inbreeding features. Number of ROH (NROH) ≥1 Mb versus the sum ROH length (SROH) ≥1 Mb per individual for each population: Africa (ESN-Esan in Nigeria, GWD- Mandinka in Gambia, LWK-Luhya in Kenya, MSL-Mende in Sierra Leone, and YRI-Yoruba in Nigeria); AP (Oman, Saudi Arabia, UAE and Yemen); East Asia (CDX-Dai in China, CHB-Han in China, CHS-Han in South China, JPT-Japanese in Japan, and KHV-Kinh in Vietnam), and Europe (GBR-British in UK, IBS-Iberians in Spain, and TSI-Toscani in Italia).
